# Heterogeneous water dynamics in the Hyaluronan-DPPC Interfaces

**DOI:** 10.1101/2025.09.24.678240

**Authors:** Anirban Paul, Jaydeb Chakrabarti

## Abstract

The synergistic interactions between hyaluronic acid (HA) and other cellular components, particularly lipids and water, play a vital role in maintaining cellular structure and function. Variations in HA concentration and chain length lead to changes in the mechanical and viscoelastic properties of diseased cells, such as those in colon cancer and osteoarthritis. Despite its biological significance, the microscopic dynamics at the HA–water and lipid bilayer interface remain poorly understood. In this study, we investigate the dynamic properties of water molecules at the interface between HA and Dipalmitoylphosphatidyl-choline (DPPC) lipid bilayer using molecular dynamics simulations. The interfacial and hydration water exhibit non-Gaussian self van Hove functions, indicating dynamic heterogeneity. The diffusivity distributions reveal contrasts in the underlying translational and rotational dynamics between the two regions. In the diffusive interface, while diffusion coefficients decrease with increasing HA concentration, they show only a weak dependence on HA chain length. However, water dynamics in the subdiffusive hydration layer are largely unaffected by the presence of HA chains.

## I. INTRODUCTION

A multi-component complex system often has dynamically heterogeneous regions^1–3^ with closely spaced relaxation time scales. The system undergoes formation of clusters of particles with similar mobilities, a situation known as dynamic heterogeneity^2,4^. In such systems, the particles get stuck in the local dynamic environment for some time and explore the entire phase space only if observed for a long enough time, exceeding the slowest relaxation in the system. The particles in the short time scale show different diffusivity of comparable magnitude depending on the local environment in which they are stuck, and manifest as dynamical heterogeneity for shorter observation times. After long enough time, however, the particles regain normal diffusive behaviour^3^.

The diffusion of particles in a fluid is, in general, given by the slope of the mean squared displacements (MSD) if the MSD is linear in time in the long-time limit. In the long-time limit, any information on the heterogeneous dynamics is lost, and hence, MSD is not a good marker for dynamic heterogeneity. Dynamic heterogeneity is signaled via the self van Hove correlation function (self-vHf), G(*ξ*, Δt), that describes the probability distribution of displacements *ξ* (translation or rotation) of the particles in time interval Δt^5–7^, the MSD being the second moment of the self-vHf^8^.

The self-vHf is experimentally measurable through scattering techniques^8^. For normal fluids with diffusive particles, the self-vHf assumes a Gaussian form whose width increases linearly with time, the slope being proportional to the diffusion coefficient D_*ξ*_ ^3,9^. In case of dynamic heterogeneity, *G*(*ξ*, Δt) is marked by deviation from Gaussian nature^3,10^. The non-Gaussian self-vHf is typically interpreted as the result of a superposition of independent Gaussian diffusive processes, where the system exhibits a distribution P(D_*ξ*_) of multiple, closely spaced diffusion coefficients^3,11^. This situation is also known as ‘Fickian yet Non-Gaussian’ dynamics^12^.

Recent studies report heterogeneous water dynamics in a host of bio-molecular systems, for instances, confined water molecules in protein-DNA complex^13^, hydration waters of lipid bilayers^14^ and intrinsically disordered proteins^15^, water molecules near organic and inorganic surfaces^1^, nano-confined water^5,16^ and so on. Aqueous suspension of Hyaluronic acid (HA), a polyanionic glycan molecule, is present in extracellular matrices of cells, synovial fluid, and other biological environments^17^. The interaction between aqueous HA suspension and lipids is shown to be crucial for various biological functions such as bio-lubrication^18,19^, water permeation through proteins^20^, and so on. The size and concentration of HA chains differ between normal and cancer cells, which is used as a colon cancer biomarker^21,22^. Moreover, these variations of HA properties lead to changes in the mechanical and viscoelastic response of the diseased cells^23–25^. Thus, the water-HA-lipid ternary system is important pedagogically as well as for biomedical applications and is yet far from being understood. In our previous study on the water-lipid bilayer-HA ternary system, we have shown the increased flexibility of the Dipalmitoylphosphatidylcholine (DPPC) lipid bilayer in the presence of HA chains^25^. We also reported the simultaneous existence of a sub-diffusive hydration water layer and diffusive water molecules in the system^26^. Water, being the ubiquitous medium for bio-molecular processes, heterogeneity in water dynamics is of utmost importance to understand^1,27^. However, heterogeneous dynamics at the interface of HA-water and lipid bilayer have not been investigated so far.

In this paper, we investigate the dynamics of water molecules at the interface of hyaluronic acid (HA) and Dipalmitoylphosphatidylcholine (DPPC) bilayer using molecular dynamics simulation. Our simulation model represents an extracellular environment near a cancer cell membrane, which is typically rich in phosphatidylcholine lipids^28–30^ and shorter HA chains^21,22^. We calculate the self van Hove functions (self-vHf) for translational dynamics, G_T_(r, Δt), and rotational dynamics, G_R_(*ϕ*, Δt). We observe non-Gaussian self-vHfs for water molecules in both interfacial and hydration layers, indicating heterogeneous dynamics. To further probe this behavior, we compute the probability distribution of diffusion coefficients from the self-vHfs^11^. The diffusivity distributions reveal distinct behaviors for translational and rotational motions in both the interfacial and hydration layer. At the interface, translational diffusivity is broadly distributed, while rotational diffusivity shows a sharp primary peak along with smaller secondary peaks. In this region, diffusion coefficients decrease with increasing HA concentration, but show only a weak dependence on HA chain length. On the contrary, in the hydration layer, translational diffusivity displays a sharp peak at low values, suggesting localized motion, whereas rotational diffusivity exhibits multiple peaks, implying dynamic exchange among angular modes. Furthermore, HA chains have minimal influence on water dynamics in the hydration layer.

## II. METHODS

### A. System preparation & simulation details

The all-atom model of a lipid bilayer of a total of 512 Dipalmitoylphosphatidylcholine (DPPC) molecules (256 in each leaflet) is constructed in a simulation box of size 12.45 nm × 12.45 nm × 13.0 nm using the CHARMM-GUI membrane builder module^31^. The bilayer is spanned in the xy plane and the normal is along the z axis. The initial structure of Hyaluronic acid is taken from RCSB PDB id 2BVK^32^ and it is modified in CHARMM GUI glycan modeler^33^ to obtain Hyaluronan monomer (HA1), pentamer(HA5) and decamer(HA10) structure. HA molecules are randomly placed above the bilayer using Packmol^34^. Sodium ions are added for electroneutrality, and additional sodium-chloride salt is added to maintain the 150 millimolar physiological salt concentration. The interaction potential is modeled using the CHARMM36 force field^35^, where the TIP3P water model is used to hydrate the system. The CHARMM36 parameters have been previously validated for HA–lipid systems^20,25,26,36^. All molecular dynamics simulations are performed using the GROMACS package^37^ with the same protocol as reported in our previous works^25,26^.

We confirm the equilibration from the saturation of area per lipid as described in ref.^25^. To analyze the interfacial dynamics, we performed multiple simulation runs, each starting from different equilibrium configurations. Trajectories were recorded at 0.5 picosecond intervals^26^, and the results were averaged over all runs.

### B. Analysis

#### 1. Translational and Rotational self van Hove function calculation

The self part of the translational van Hove function, G_T_(r,Δt) is given by the following formula^5^:

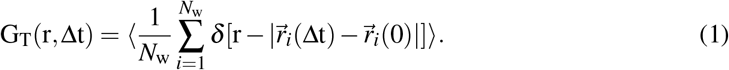

Here *N*_w_ indicates the number of water molecules that remain at the interface throughout the time interval of Δt. Here, r is displacement in two dimensions (in the lateral plane of the bilayer).

The rotational self van Hove function is defined^7,38^ as

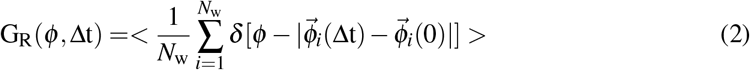

Here 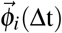 is the displacement of i’th molecule in rotational space where 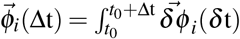. The magnitude of 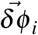 is given by 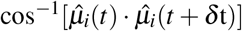 and its direction is given by 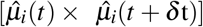 where, 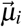 is the dipole moment of the i’th water moleucle^5,38^. 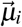 is defined as the straight line joining the oxygen atom and the center of mass of the two hydrogen atoms of the i’th water molecule. We take *δ* t = 0.5 ps. Both G_T_(r,Δt) and G_R_(*ϕ*, Δt) are calculated using unfolded trajectory.

#### 2. Fitting of G_T_(r,Δt) and G_R_(ϕ, Δt) data

To determine whether G_T_(r,Δt) has a Gaussian or exponential tail following the central Gaussian beyond a crossover length r_c_, we follow the method prescribed in previous literature^5,9,10^. First, we fit ln G_T_(r, Δt) as a double-Gaussians: for *r* < *r*_*c*_, 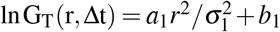 (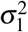 is the width of the central Gaussian), and for *r* > *r*_*c*_, 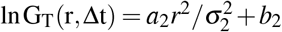 (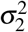 is the width of the Gaussian tail). We minimize 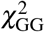 with respect to r_c_ and the fitting parameters. Next, we fit a Gaussian-exponential function: for *r* < *r*_*c*_, 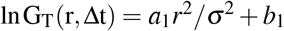 and for *r* > *r*_*c*_, ln G_T_(r, Δt) = *a*_2_*r*/*λ* + *b*_2_, where *λ* is the decay constant of the exponential tail. Again, we minimize 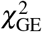 with respect to r_c_ and the fitting parameters. The fit with the smaller of 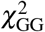 and 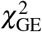 is chosen as the best functional fit.

The same procedure is also applied to G_R_(*ϕ*, Δt) to extract the tail behavior and crossover angle *ϕ*_*c*_. The exponent of the time dependence of r_c_, *ϕ*_*c*_, *σ*^2^, and *λ* is obtained by averaging and estimating the error bars over three equilibrium trajectories. The error bars are computed by dividing the standard deviation of the quantity by the square root of the number of samples.

#### 3. Distribution of diffusivity using Lucy’s de-convolution method

We compute the normalized probability distribution of translational diffusivity P(D_T_) from tvhf G(r, Δt) following the method mentioned in Ref.^11,39^. The relation between P(D_T_) and G(r, Δt) is given by^3,11^

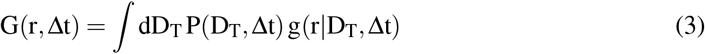

where 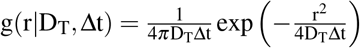

We calculate P(D_T_) iteratively from G(r, Δt) using Lucy’s deconvolution method^11,39^:

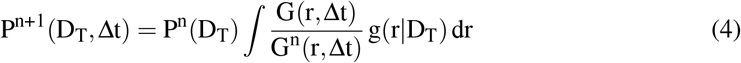

Here P^n^(D_T_) is the distribution of diffusivity obtained at the n’th iteration. The initial distribution is taken as 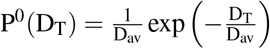. We follow the same method to compute the probability distribution of rotational diffusivity P(D_R_) as well. For rotation, we consider 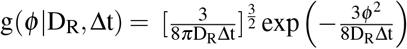 ^38,40^.

## III. RESULTS

The studies are carried out without HA chains, 10, 30, and 50 HA pentamers (HA5) (n_HA5_= 0,10, 30, and 50 respectively); and HA molecules of different chain lengths but same monomer concentration: 150 HA monomer (HA1), 30 HA pentamer (HA5), and 15 HA decamer (HA10) (N = 1, 5, and 10, respectively). Figure 1(a), 1(b) and 1(c) show the density profiles of water, phosphorus atoms of the bilayer, and HA center of mass (com), along the bilayer normal (z-axis) and with respect to the origin at the center of the bilayer, for HA free case (n_HA5_=0), n_HA5_=50 and N=10, respectively. For the HA-free case in Figure 1(a), the two peaks of the phosphorus density profile correspond to the phosphorus atoms from the upper and lower leaflets of the bilayer. The water density profile is zero inside the bilayer because of its hydrophobic core and increases to the bulk density far away from the bilayer, forming an interfacial region. The water layer within 5Å distance from the bilayer peaks in the interface is the hydration water interacting directly with the lipid head groups. Figure 1(b) and 1(c) show that the peak of the density profile of HA is 15Å away from the peak of the phosphorus atoms, approximately where the density of bulk water sets in. Thus, the HA molecules interact with the diffusive interfacial water molecules but not directly with the bilayer. The formation of water-bilayer and water-HA interfaces is in agreement with our previous reports^25^. Earlier, we showed that the water molecules within the hydration layer have a residence time *τ*_s_ = 10 ps and are subdiffusive. The water molecules beyond the hydration layer are diffusive, which we designate as diffusive interface^26^.

**FIG. 1.**
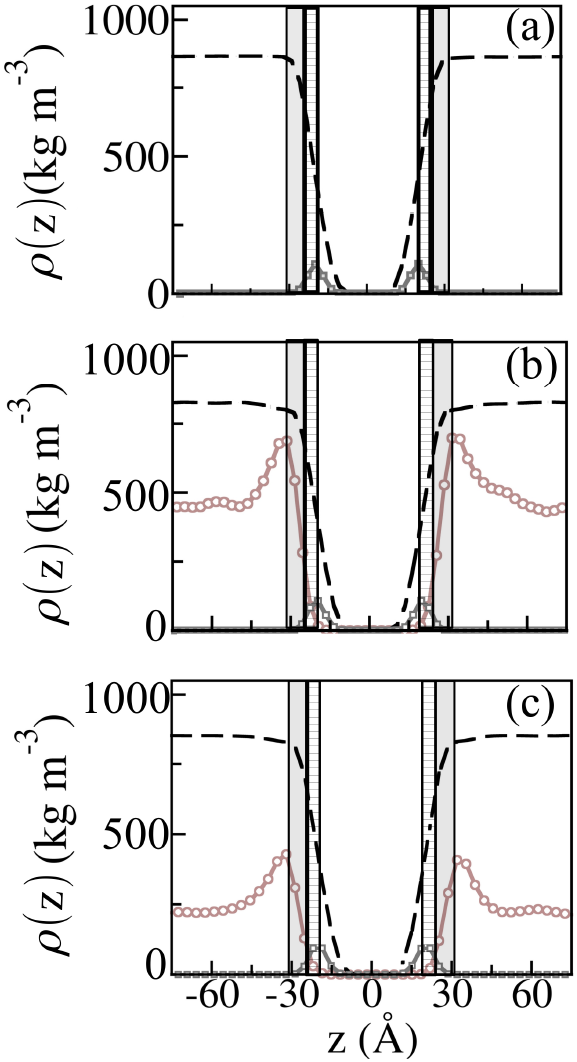
Density profile of phosphorus atoms of the lipid bilayer (*dotted gray*), water (*dashed black* line), and HA com (*dotted brown* line) along the bilayer normal for (a) n_HA5_=0, (b) n_HA5_=50, and (c) N=10. The hydration layer and interfacial layer is shown by the dashed and shaded regions, respectively. HA density profile is scaled up with a factor of 5.0 for better visualization.

We investigate the dynamics of the water molecules at the interface by computing the self van Hove function (self-vHf) for translational dynamics G_T_(r,Δt) and rotational dynamics G_R_(*ϕ*, Δt). Here, r is the distance in two dimensions traveled by the water molecules in the lateral plane of the bilayer, and *ϕ* is the rotational displacement of the water dipole vectors in the time interval Δt^38^. The self-vHfs are calculated while the water molecules reside in the interfacial or hydration layer. Hence Δt is restricted within the residence time of water in a particular region. We also calculate the probability distribution of diffusion coefficients from the self-vHf data^11,39^.

### A. Dynamics of diffusive interfacial water layer

First, we consider the water molecules in the diffusive interface. We define the time interval scaled by the residence time of the hydration water molecules, Δt* = Δt/*τ*_s_. We consider G_T_(r, Δt*) and G_R_(*ϕ*, Δt*) for the diffusive interfacial water molecules for Δt* = 2.0 for HA free case. Note, the mean residence time of water at the diffusive interface is 20 ps^26^. Figure 2(a) shows lnG_T_(r,Δt*) vs r data for Δt*= 2.0. We find that G_T_(r, Δt*) is double Gaussian in nature. The crossover length r_c_ between the two Gaussian curves, shown in SI Figure S1(a), shows a sublinear dependence, indicating that the two distinct Gaussians persist up to the residence time of the interfacial water. The double Gaussian G_T_(r, Δt*) indicates the presence of two diffusion coefficients at the interface. The widths of the two Gaussians, 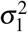 and 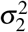, as shown in SI Figure S1(b) and S1(c), respectively, grow linearly with time, consistent with the diffusive dynamics reported earlier^26^. From the slope of 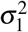 vs Δt and 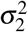 vs Δt data, we compute two diffusion coefficients (*σ*^2^ = 4D_T_Δt, see *section B*.*3* of Methods), 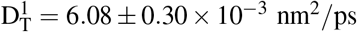 and 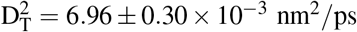. The values are close, which is a signature of the coexistence of dynamically heterogeneous regions with closely spaced relaxation times. Figure 2(b) shows the underlying distribution P(D_T_) derived from G_T_(r, Δt*) using Lucy’s deconvolution method^11,39^. P(D_T_) shows a broad distribution of D_T_. The average diffusion constant ⟨D_T_⟩(= 6.40 ± 0.07 × 10^−3^ nm^2^/ps), computed from P(D_T_) is close to the mean value of 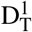 and 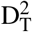. ⟨D_T_⟩ is also comparable to the translational diffusion coefficient obtained from MSD data in Ref^26^. This further demonstrates that MSD data in the long-time limit is averaged over dynamically heterogeneous regions.

**FIG. 2.**
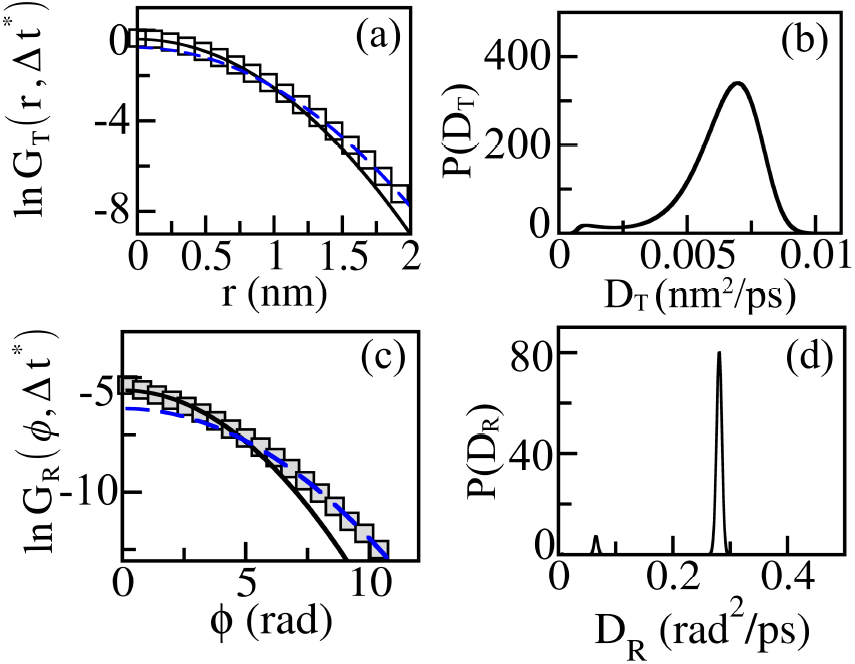
(a) lnG_T_(r,Δt*) vs r for n_HA5_ = 0 for Δt*=2.0 in the diffusive interface. (b) Normalized probability distribution of translational diffusivities P(D_T_) calculated from G_T_(r,Δt*) (c) lnG_R_(*ϕ*, Δt*) vs *ϕ* for n_HA5_ = 0 for Δt*=2.0 in the diffusive interface. (d) Normalized probability distribution of rotational diffusivities P(D_R_) calculated from G_R_(*ϕ*,Δt*). In the self-vHf plots, *Solid* lines show the best fitted central Gaussian, where *broken* lines imply the best fitted Gaussian tail.

The rotational self-vHF G_R_(*ϕ*, Δt*) for Δt*=2.0 in Figure 2(c) also shows a double Gaussian form. The non-Gaussian G_R_(*ϕ*, Δt*) indicates dynamic heterogeneity in rotation for the interfacial waters in the diffusive layer. 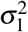 and 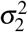 for the double Gaussian profile grow linearly with time as shown in SI Figure S2. Two diffusion coefficients calculated from the slope of the time dependence of 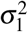 and 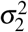 (for rotation, 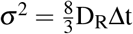, see *section B*.*3* of Methods), 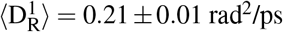 and 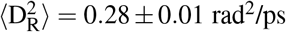, show close values as in the case of translational motion. Figure 2(d) show P(D_R_) at Δt*=2.0. We observe a sharp peak at ⟨D_R_⟩=0.27 rad^2^/ps with a very small peak at very small D_R_. We note that the sharper peak is located near the value 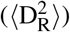 obtained from the slope of linear time dependencies of 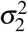 and also agrees well with the previous estimate from MSD data^26^.

Now we describe how the translational and rotational dynamics of the diffusive water layer respond to the presence of HA chains. First, we consider the n_HA5_ dependence. We find double Gaussian G_T_(r, Δt*) at Δt* = 2.0 (SI Figure S3(a)-(c)) similar to HA-free case. The variation of r_c_ with time follows a sublinear dependence (SI Figure S4(a)), while the widths of the two Gaussians 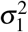 and 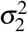 vary linearly with time (SI Figure S4(b)-(c)) as observed in the HA-free case. The underlying distribution of diffusion coefficients P(D_T_) for different n_HA5_ is shown in Figure 3(a). Figure 3(b) shows the average translational diffusion coefficient, ⟨D_T_⟩, for different values of n_HA5_. We observe that ⟨D_T_⟩ decreases linearly as n_HA5_ increases.

**FIG. 3.**
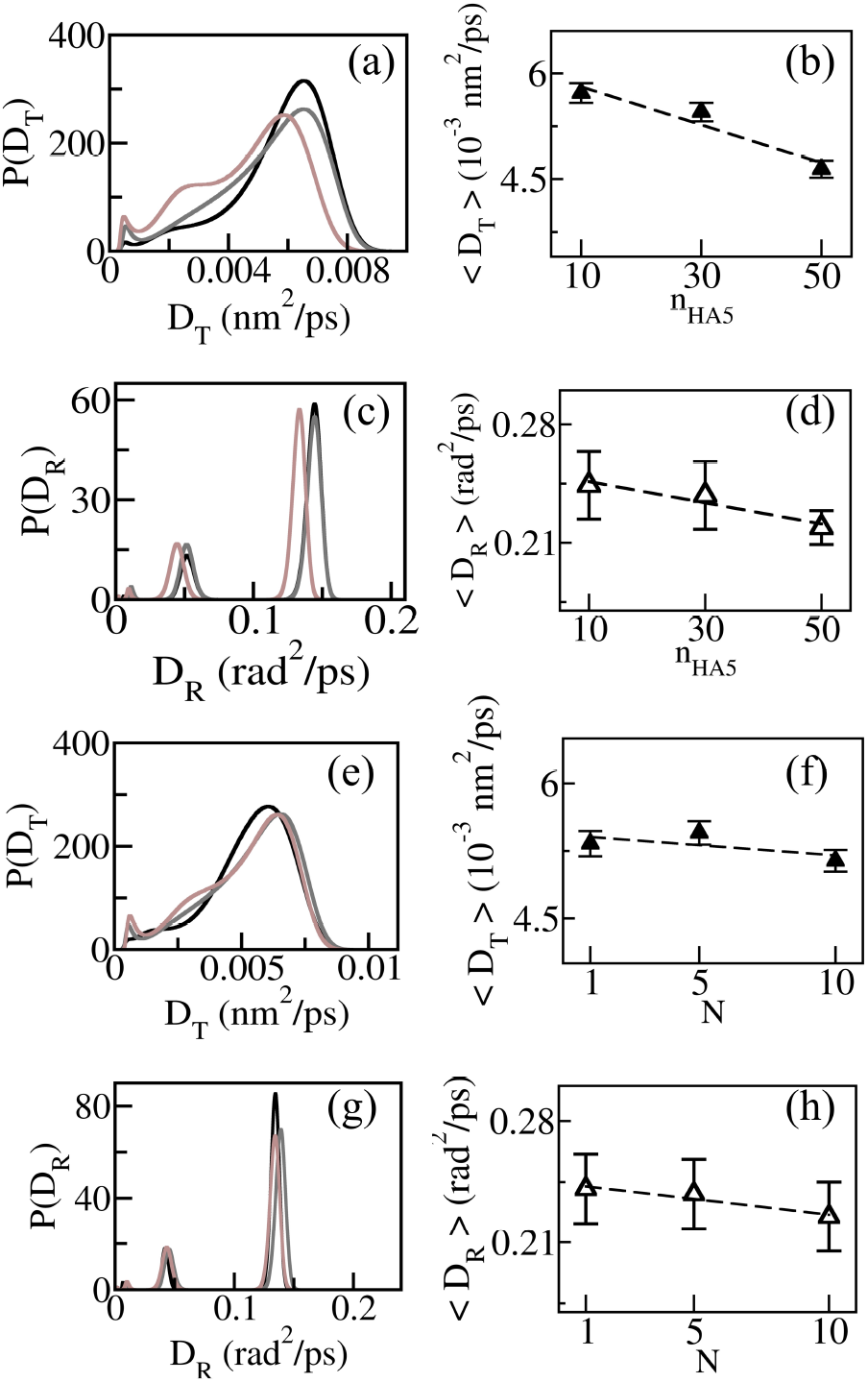
(a) Normalized probability distribution of diffusivity P(D_T_) of the interfacial water for n_HA5_=10(*black*), 30 (*gray*) and 50 (*brown*) for Δt*=2.0. (b) ⟨D_T_⟩ for different n_HA5_. (c) P(D_R_) of the interfacial water for n_HA5_=10(*black*), 30 (*gray*) and 50 (*brown*) for Δt*=2.0. (d) ⟨D_R_⟩ for different n_HA5_. (e) P(D_T_) of the interfacial water for N=1 (*black*), N=5 (*gray*) and N=10 (*brown*) (f) ⟨D_T_⟩ for different N. (g) P(D_R_) of the interfacial water for N=1 (*black*), N=5 (*gray*) and N=10 (*brown*) (h) ⟨D_R_⟩ for different N.

The G_R_(*ϕ*, Δt*) for the diffusive interfacial water at Δt* = 2.0 for varying n_HA5_ are shown in SI Figure S5(a)-(c)). We find that G_R_(*ϕ*, Δt*) is double Gaussian in nature as in the HA-free case. For all n_HA5_, the P(D_R_) distributions exhibit a prominent peak around ⟨D_R_⟩, along with smaller peaks at lower D_R_ values. We note that the smaller peaks appear only in the presence of HA chains, suggesting the existence of water molecules with multiple rotational modes. Figure 3(d) shows that ⟨D_R_⟩ decreases with n_HA5_. Hence, both the translation and rotation of the water molecules become increasingly sluggish with increasing HA concentration, which is in agreement with our previous report^26^.

G_T_(r,Δt)(SI Figure S6(a)-(c)) and G_R_(*ϕ*, Δt) (SI Figure S7(a)-(c)) for interfacial water molecules for different chain size N behave similarly as in the case of different HA concentration. However, P(D_T_) and P(D_R_), unlike the dependence on n_HA5_, show insensitivity to N as shown in Figure 3(e) and 3(g) respectively. ⟨D_T_⟩ and ⟨D_R_⟩ from the respective distributions also show weak dependence on N, as described in Figure 3(f) and 3(h), which is in agreement with our earlier findings^26^.

### B. Dynamics of Hydration water

Next, we examine the dynamic property of the hydration water molecules where the translational and rotational dynamics are observed to be subdiffusive^26^. Let us first consider the HA-free case. We compute the translational self-vHf G_T_(r,Δt*) for two observation times, namely Δt*=0.5 and 1.0, shown in semi-logscale in figure 4(a). G_T_(r,Δt*) shows a central Gaussian followed by an exponential tail in both cases. Such exponential tails in self-vHf have been reported earlier in different soft-matter systems such as liposomal solution^3^, colloids^9,41^, glasses^42^, and so on. This indicates the dynamic heterogeneity of the hydration water, namely, coexisting regions with different diffusion coefficients within a narrow range of values. We find in Figure 4(b) that the cross-over distance between the central Gaussian and exponential tail, r_c_ show sublinear dependence. The sublinear dependence of r_c_ means that the dynamic heterogeneity persists till the water molecules reside in the hydration layer. We also show sublinear time variation of the width of the central Gaussian *σ*^2^ and decay length of the exponential tail *λ* in the inset of Figure 4(b). The time dependence of *σ*^2^ is consistent with the sub-diffusive behaviour observed earlier from the MSD data^26^. This is in contrast with the linear dependence of the Gaussian width parameters observed in the case of diffusive interfacial water molecules. Now, we compute the normalized probability distribution of diffusivity P(D_T_) from G_T_(r,Δt*)^11,39^. We show data for Δt*=0.5 and 1.0 in Figure 4(b). A sharp peak is observed in P(D_T_) at a very small D_T_ value with a long tail for both time intervals. The sharp peak at a very small D_T_ value corresponds to dynamically arrested water molecules near the bilayer. The long tail indicates highly heterogeneous dynamics at the hydration layer.

**FIG. 4.**
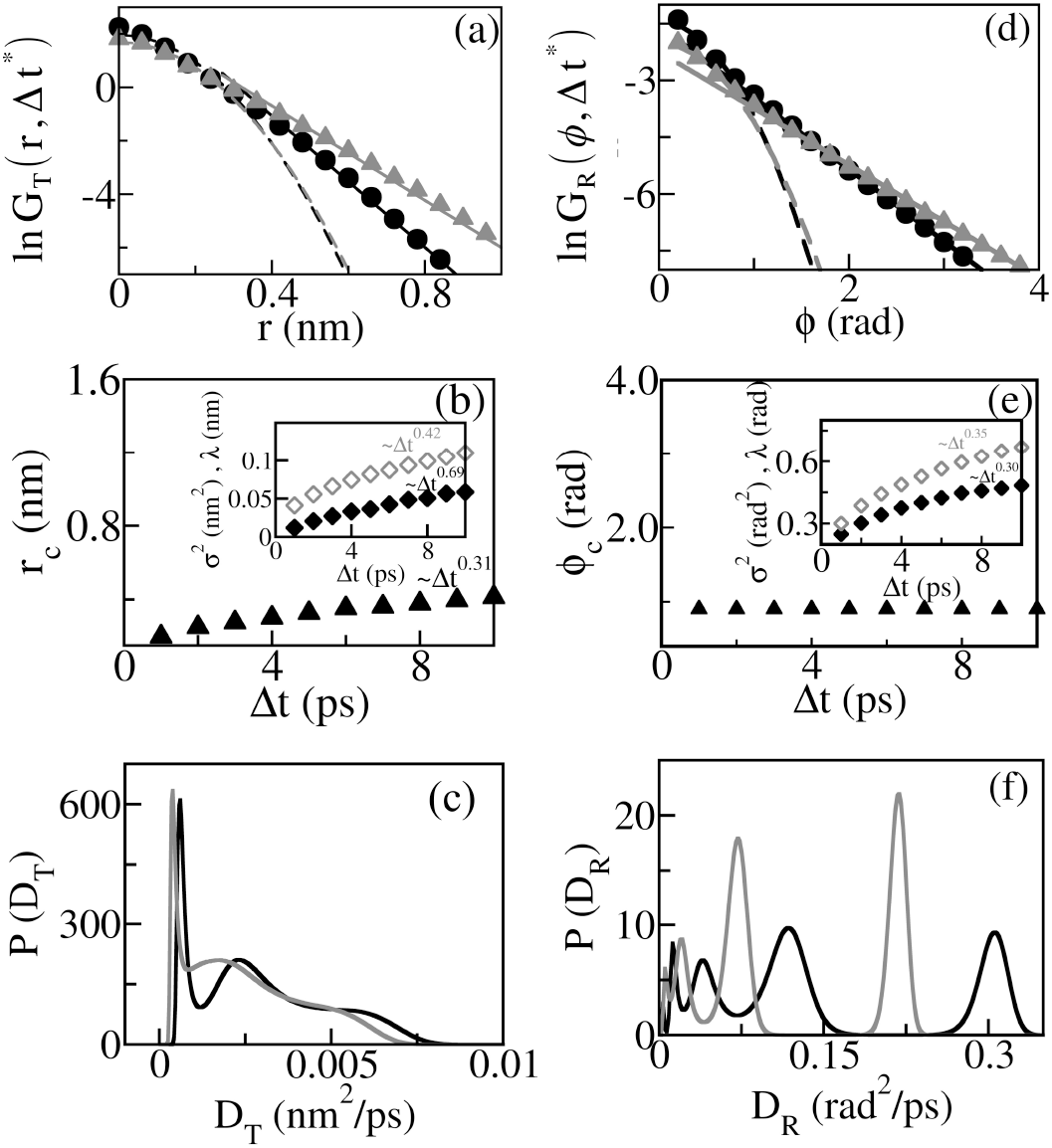
(a) lnG_T_(r,Δt*) vs Δt plot for n_HA5_ = 0 for Δt*= 0.5 (*circles*) and 1.0 (*triangles*) in the hydration layer. *Broken* and *solid* lines show the fitted central Gaussian, and the exponential tails. (b) r_c_ vs Δt. r_c_ ~ Δt^0.3^ inset: time variation of *σ*^2^ (*closed* symbols) and *λ* (*open* symbols). *σ*^2^ ~ t^0.69^, *λ* ~ t^0.42^ (c) P(D_T_) for Δt*= 0.5 (*black*) and 1.0 (*gray*)(d) lnG_R_(*ϕ*, Δt*) vs Δt plot for n_HA5_ = 0 for Δt*= 0.5 (*circles*) and 1.0 (*triangles*) in the hydration layer. *Broken* lines show the fitted central Gaussian, whereas *solid* lines imply the exponential Gaussian tails. (e) *ϕ*_*c*_ vs Δt. inset: time variation of *σ*^2^ (*closed* symbols) and *λ* (*open* symbols). *σ*^2^ ~ t^0.30^, *λ* ~ t^0.35^ (f) P(D_R_) for Δt*= 0.5 (*black*) and 1.0 (*gray*)

The rotational self-vHf G_R_(*ϕ*, Δt*) data in figure 4(d) show long exponential tails for both time intervals of Δt*= 0.5 and 1.0. The crossover angle *ϕ*_*c*_ remains unchanged with time, as shown in Figure 4(e). In the inset of Figure 4(e), we show the time variation of *σ*^2^ of the central Gaussian and *λ* of the exponential tail. We find that both *σ*^2^ and *λ* vary sub-linearly, qualitatively similar to the translation case, and consistent with the sub-diffusive rotational motion of hydration water^26^. Figure 4(f) shows the probability distributions of rotational diffusion constants P(D_R_). P(D_R_) exhibits multiple peaks of similar amplitude, unlike the diffusive water layer. The peaks at extremely small D_R_ correspond to the water molecules that are stuck to the bilayer with little rotational freedom. However, multiple peaks imply the coexistence of slow and fast species of water molecules in this region.

Next, we show lnG_T_(r, Δt*) vs r data for n_HA5_=10, 30 and 50 in SI Figure S8(a)-(c). In the hydration layer, G_T_(r,Δt*) exhibits a central Gaussian with exponential tails similar to the HA free case. Moreover, the variations of r_c_ (SI Figure S9(a)), *σ*^2^, and *λ* (SI Figure S9(b)) with time show power law dependencies as in the HA-free case. P(D_T_) for the two different time intervals are shown in Figure 5(a) and 5(b). We observe a sharp peak at a small D_T_ value, and a wide range of diffusivity contributes to the distribution. While HA concentration has a negligible effect on the tail of the distribution at Δt*= 0.5, the peak of P(D_T_) enhances at the small D_T_ value for n_HA5_=50 at Δt*=1.0. It shows that the water molecules of the lipid hydration layer slow down at high HA concentration.

**FIG. 5.**
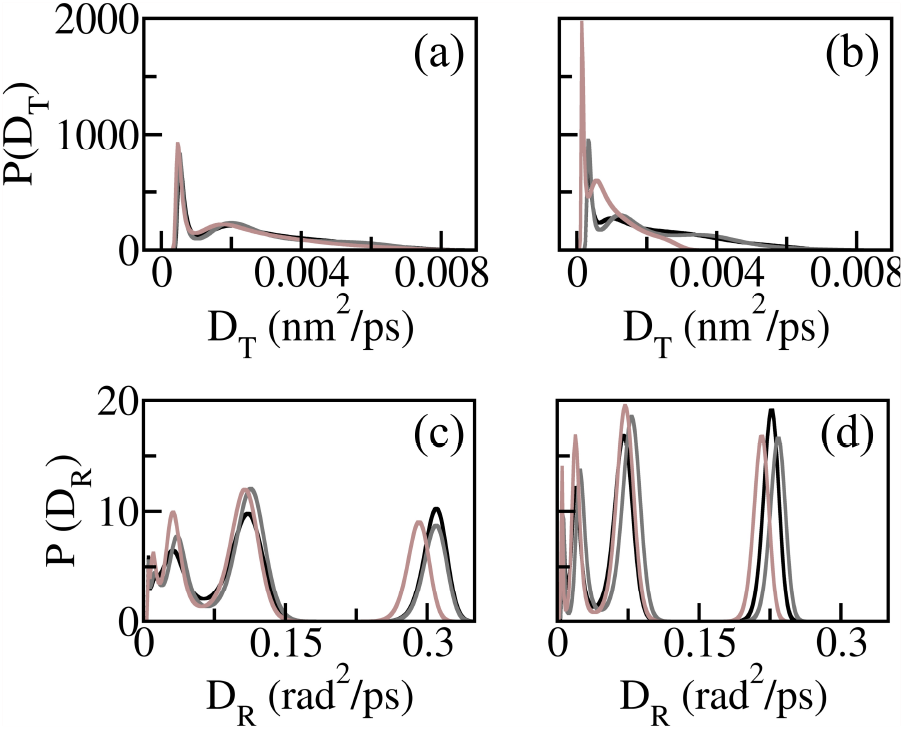
Normalized probability distribution of translational diffusivity P(D_T_) of the hydration water for n_HA5_=10(*black*), 30 (*gray*) and 50 (*brown*) for (a) Δt*= 0.5 and (b) 1.0. Normalized probability distribution of rotational diffusivity P(D_R_) of the hydration water for n_HA5_=10(*black*), 30 (*gray*) and 50 (*brown*) for (c) Δt*= 0.5 and (d) 1.0.

The rotational self-vHf G_R_(*ϕ*, Δt*) for n_HA5_=10,30, and 50 (shown in SI Figure S10(a)-(c)) in the hydration layer behave similarly to the HA-free cases. In all three cases, we observe an exponential tail following the central Gaussian. The crossover angle *ϕ*_*c*_ does not change with time (SI Figure S11(a)). *σ*^2^, and *λ* (SI Figure S11(b)) show power law dependencies as in HA-free case. P(D_R_) curves, shown in Figures 5(c) and 5(d), exhibit multiple peaks, implying multiple modes of rotational diffusion.

Next, we show lnG_T_(r,Δt*) vs Δt* data for different HA chain sizes, N=1,5 and 10 in SI Figure S12(a)-(c) for hydration water. For all cases, G_T_(r,Δt*) show a central Gaussian with an exponential tail. r_c_ shows sublinear dependence on time (SI Figure S13(a)). Hence, the heterogeneity persists till the residence time of water in the hydration layer. *σ*^2^ and *λ* show sublinear dependence with time (SI Figure S13(b)). P(D_T_) data in Figures 6(a) and 6(b) show peaks at a very small value of D_T_, and contribution from a wide range of D_T_ values is noted, suggesting dynamic heterogeneity. We find that P(D_T_) remains similar for all values of N.

**FIG. 6.**
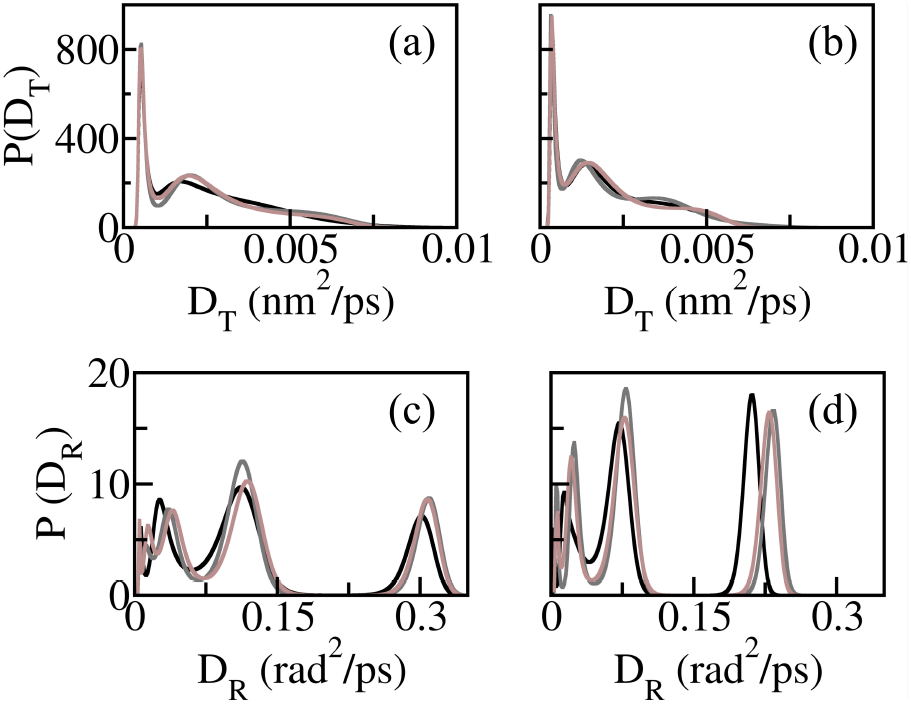
Distribution of diffusivity P(D_T_) for the hydration water for N=1(*black*), 5 (*gray*) and 10 (*brown*) for (a) Δt*=0.5 and (b) Δt*=1.0. P(D_*R*_) for the hydration water for N=1(*black*), 5 (*gray*) and 10 (*brown*) for (c) Δt*=0.5 and (d) Δt*=1.0.

The rotational self-vHf G_R_(*ϕ*, Δt*) (Figure SI Figure 14(a)-(c)) for the hydration water molecules has a central Gaussian followed by an exponential tail for all N. We find that the crossover angle *ϕ*_*c*_ remains unchanged with time (SI Figure S15(a)). The width *σ*^2^ of the central Gaussian and the decay constant *λ* of the exponential tail show sublinear power-law time dependence (SI Figure S15(b)). P(D_R_), shown in Figure 6(c) and 6(d), have multiple peaks, implying water molecules with different rotational diffusivity. The P(D_R_) is almost the same for all N values, as same as P(D_T_).

The heterogeneity in the hydration water dynamics near the lipid bilayer is consistent with earlier reports^14^. Furthermore, here we show that the distributions of translational and rotational diffusion coefficients of the hydration water remain nearly unchanged for different values of n_HA5_ and N. Thus, the presence of HA chains does not significantly affect the dynamics of hydration water, which is likely to be governed more by the bilayer head groups than by the HA chains.

Next, we extract both the translational MSD, ⟨r^2^⟩ and the rotational MSD, ⟨*ϕ*^2^⟩ for the hydration water from G_T_(r, Δt) and G_R_(*ϕ*, Δt). For that, we compute the second moment of G_T_(r, Δt) and G_R_(*ϕ*, Δt) for different Δt up to residence time *τ*_*s*_. Figure 7(a) and (b) present the ⟨r^2^⟩ for various values of n_HA5_ and N respectively. Figure 7(c) and (d) depict the ⟨*ϕ*^2^⟩ for different n_HA5_ and N respectively. We find that water molecules in the hydration layer exhibit subdiffusive behavior in both translational and rotational dynamics for different n_HA5_ and N. This is consistent with our earlier report, where we demonstrate the subdiffusive nature by directly computing the MSD^26^.

**FIG. 7.**
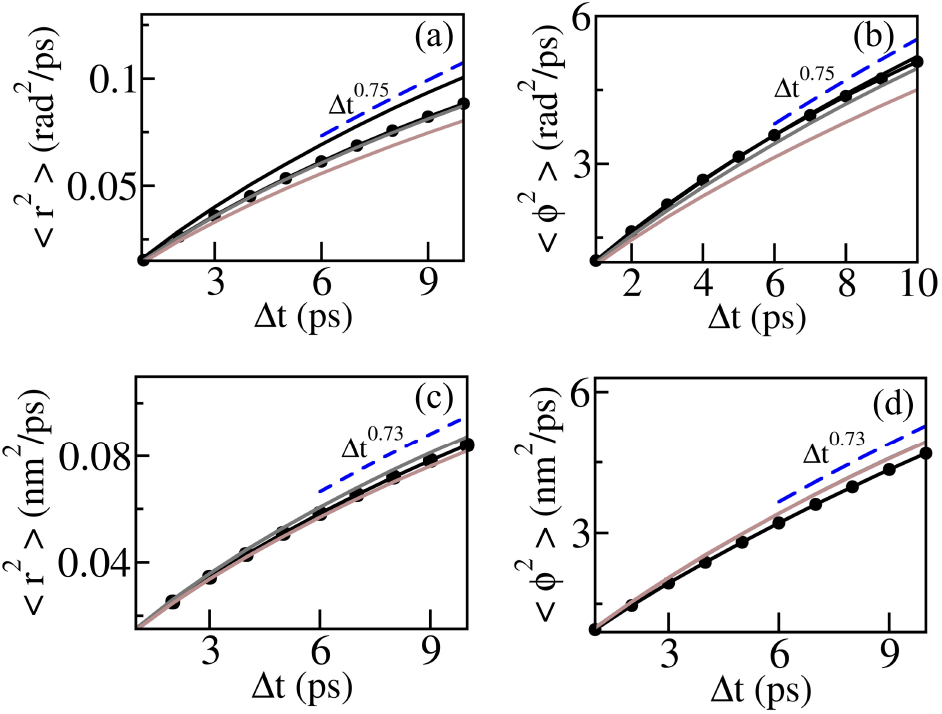
(a) Translational MSD of water ⟨r^2^⟩ and (b) Rotational MSD of water ⟨*ϕ*^2^⟩ in hydration layer for n_HA5_=0 (*solid black line*), 10 (*dotted black line*), 30 (*gray line*) and 50 (*brown line*). (c) ⟨r^2^⟩ and (d) ⟨*ϕ*^2^⟩ in the hydration layer for N=1 (*dotted black line*), 5 (*gray line*) and 10 (*brown line*).

Despite being subdiffusive, P(D_T_) and P(D_R_) reveal distinctions between the translational and rotational dynamics of the hydration water. For subdiffusive translation of the hydration waters, P(D_T_) exhibits a sharp peak near vanishing diffusivity values, indicative of localized or trapped motion of water molecules. In contrast, for subdiffusive rotation, P(D_R_) displays multiple peaks, which suggests possible dynamic exchange among discrete angular modes.

## IV. CONCLUSION

In summary, we present a systematic study of the dynamic heterogeneity of water molecules at the HA-water and DPPC interface, focusing on how the HA concentration and HA chain size influence the dynamics. The interfacial and hydration water at the HA-water and DPPC interface exhibit dynamic heterogeneity, characterized by non-Gaussian self van Hove functions. Furthermore, the diffusivity distributions provide a detailed understanding of the underlying dynamics. In the interfacial layer, translational diffusivity exhibits a broad distribution, whereas rotational diffusivity displays a dominant peak with minor secondary modes. Increased HA concentration decreases both translational and rotational diffusion at the interface, whereas HA chain length has minimal influence. In the hydration layer, translational diffusivity is sharply peaked at very low translational diffusivity, indicating highly restricted motion, while its rotational distributions are multimodal. Importantly, HA chains have minimal impact on hydration-layer dynamics. Our study provides insights into the interfacial dynamics of HA-lipid complexes, which could help in their spectroscopic characterization, with potential implications in many biomedical applications.

## AUTHOR CONTRIBUTIONS

**Anirban Paul:** conceptualization, data curation, formal analysis, methodology, validation, visualization, software, investigation, writing – original draft, writing – review and editing. **J. Chakrabarti:** conceptualization, methodology, validation, project administration, resources, supervision, writing – review and editing.

## DATA AVAILABILITY STATEMENT

The data that support the findings of this study are available within the article and its supplementary material.

## ACKNOWLEDGMENTS

The authors thank the Technical Research Centre at S. N. Bose National Centre for Basic Sciences, Kolkata for its computational facilities. A.P acknowledges financial support from Council of Scientific and Industrial Research (CSIR), India [File No. 09/575(0134)/2020-EMR-I] and SNBNCBS for ‘Bridge Fellowship’.

## REFERENCES

1 Sander Pronk, Erik Lindahl, and Peter M Kasson. Dynamic heterogeneity controls diffusion and viscosity near biological interfaces. Nature communications, 5(1):3034, 2014.

2 Juan P Garrahan. Dynamic heterogeneity comes to life. Proceedings of the national academy of sciences, 108(12):4701–4702, 2011.

3 Bo Wang, James Kuo, Sung Chul Bae, and Steve Granick. When brownian diffusion is not gaussian. Nature materials, 11(6):481–485, 2012.

4 Indrajit Tah, Anoop Mutneja, and Smarajit Karmakar. Understanding slow and heterogeneous dynamics in model supercooled glass-forming liquids. ACS omega, 6(11):7229–7239, 2021.

5 E Tendong, T Saha Dasgupta, and J Chakrabarti. Dynamics of water trapped in transition metal oxide-graphene nano-confinement. Journal of Physics: Condensed Matter, 32(32):325101, 2020.

6 Marco G Mazza, Nicolas Giovambattista, H Eugene Stanley, and Francis W Starr. Connection of translational and rotational dynamical heterogeneities with the breakdown of the stokes-einstein and stokes-einstein-debye relations in water. Physical Review E, 76(3):031203, 2007.

7 Qiang Zhang, TianMin Wu, Chen Chen, Shaul Mukamel, and Wei Zhuang. Molecular mechanism of water reorientational slowing down in concentrated ionic solutions. Proceedings of the National Academy of Sciences, 114(38):10023–10028, 2017.

8 Jean-Pierre Hansen and Ian Ranald McDonald. Theory of simple liquids: with applications to soft matter. Academic press, 2013.

9 Suman Dutta and J Chakrabarti. Anomalous dynamical responses in a driven system. Europhysics Letters, 116(3):38001, 2016.

10 Sutapa Dutta, Mahua Ghosh, Rahul Karmakar, and J Chakrabarti. Dynamic signature of ligand binding over a protein surface. Physical Review E, 100(6):062411, 2019.

11 Shiladitya Sengupta and Smarajit Karmakar. Distribution of diffusion constants and stokes-einstein violation in supercooled liquids. The Journal of chemical physics, 140(22), 2014.

12 Alejandro Cuetos, Neftalí Morillo, and Alessandro Patti. Fickian yet non-gaussian diffusion is not ubiquitous in soft matter. Physical Review E, 98(4):042129, 2018.

13 Song-Ho Chong and Sihyun Ham. Anomalous dynamics of water confined in protein–protein and protein–dna interfaces. The journal of physical chemistry letters, 7(19):3967–3972, 2016.

14 Abhinav Srivastava, Sheeba Malik, and Ananya Debnath. Heterogeneity in structure and dynamics of water near bilayers using tip3p and tip4p/2005 water models. Chemical Physics, 525:110396, 2019.

15 Souvik Mondal, Krishna Prasad Ghanta, and Sanjoy Bandyopadhyay. Dynamic heterogeneity at the interface of an intrinsically disordered peptide. Journal of Chemical Information and Modeling, 62(8):1942–1955, 2022.

16 Mehdi Karzar Jeddi and Santiago Romero-Vargas Castrillon. Dynamics of water monolayers confined by chemically heterogeneous surfaces: Observation of surface-induced anisotropic diffusion. The Journal of Physical Chemistry B, 121(41):9666–9675, 2017.

17 Grigorij Kogan, Ladislav Šoltés, Robert Stern, Jürgen Schiller, and Raniero Mendichi. Hyaluronic acid: its function and degradation in in vivo systems. Studies in natural products chemistry, 34:789–882, 2008.

18 Min Wang, Chao Liu, Esben Thormann, and Andra Dedinaite. Hyaluronan and phospholipid association in biolubrication. Biomacromolecules, 14(12):4198–4206, 2013.

19 Chao Liu, Min Wang, Junxue An, Esben Thormann, and Andra Dėdinaitė. Hyaluronan and phospholipids in boundary lubrication. Soft Matter, 8(40):10241–10244, 2012.

20 Hong Zhang, Wensheng Cai, and Xueguang Shao. Regulation of aquaporin-3 water permeability by hyaluronan. Physical Chemistry Chemical Physics, 23(45):25706–25711, 2021.

21 Debashish Paul, Anuradha Roy, Arpita Nandy, Brateen Datta, Prateeka Borar, Samir Kumar Pal, Dulal Senapati, and Tatini Rakshit. Identification of biomarker hyaluronan on colon cancer extracellular vesicles using correlative afm and spectroscopy. The Journal of Physical Chemistry Letters, 11(14):5569–5576, 2020.

22 Guoliang Zhang, Renquan Lu, Man Wu, Yiwen Liu, Yiqing He, Jing Xu, Cuixia Yang, Yan Du, and Feng Gao. Colorectal cancer-associated á6ákda hyaluronan serves as a novel biomarker for cancer progression and metastasis. The FEBS Journal, 286(16):3148–3163, 2019.

23 Zhang Liu, Weifeng Lin, Yaxun Fan, Nir Kampf, Yilin Wang, and Jacob Klein. Effects of hyaluronan molecular weight on the lubrication of cartilage-emulating boundary layers. Biomacromolecules, 21(10):4345–4354, 2020.

24 Zhixiang Cai, Hongbin Zhang, Yue Wei, Min Wu, and Ailing Fu. Shear-thinning hyaluronan-based fluid hydrogels to modulate viscoelastic properties of osteoarthritis synovial fluids. Biomaterials science, 7(8):3143–3157, 2019.

25 Debashish Paul, Anirban Paul, Dipanjan Mukherjee, Saroj Saroj, Manorama Ghosal, Suchetan Pal, Dulal Senapati, Jaydeb Chakrabarti, Samir Kumar Pal, and Tatini Rakshit. A mechanoelastic glimpse on hyaluronan-coated extracellular vesicles. The Journal of Physical Chemistry Letters, 13(36):8564–8572, 2022.

26 Anirban Paul and Jaydeb Chakrabarti. Dynamics of an aqueous suspension of short hyaluronic acid chains near a dppc bilayer. Phys. Chem. Chem. Phys., 26:20440–20449, 2024.

27 Damien Laage, Thomas Elsaesser, and James T Hynes. Water dynamics in the hydration shells of biomolecules. Chemical Reviews, 117(16):10694–10725, 2017.

28 Wasinee Khuntawee, Rawiporn Amornloetwattana, Wanwipa Vongsangnak, Katawut Namdee, Teerapong Yata, Mikko Karttunen, and Jirasak Wong-Ekkabut. In silico and in vitro design of cordycepin encapsulation in liposomes for colon cancer treatment. RSC advances, 11(15):8475– 8484, 2021.

29 AB Hendrich and K Michalak. Lipids as a target for drugs modulating multidrug resistance of cancer cells. Current drug targets, 4(1):23–30, 2003.

30 Thomas E Merchant, Pamela M Diamantis, Gregory Lauwers, Toni Haida, John N Kasimos, Jose Guillem, Thomas Glonek, and Bruce D Minsky. Characterization of malignant colon tumors with 31p nuclear magnetic resonance phospholipid and phosphatic metabolite profiles. Cancer, 76(10):1715–1723, 1995.

31 Emilia L Wu, Xi Cheng, Sunhwan Jo, Huan Rui, Kevin C Song, Eder M Dávila-Contreras, Yifei Qi, Jumin Lee, Viviana Monje-Galvan, Richard M Venable, et al. Charmm-gui membrane builder toward realistic biological membrane simulations, 2014.

32 Andrew Almond, Paul L DeAngelis, and Charles D Blundell. Hyaluronan: the local solution conformation determined by nmr and computer modeling is close to a contracted left-handed 4-fold helix. Journal of molecular biology, 358(5):1256–1269, 2006.

33 Sang-Jun Park, Jumin Lee, Yifei Qi, Nathan R Kern, Hui Sun Lee, Sunhwan Jo, InSuk Joung, Keehyung Joo, Jooyoung Lee, and Wonpil Im. Charmm-gui glycan modeler for modeling and simulation of carbohydrates and glycoconjugates. Glycobiology, 29(4):320–331, 2019.

34 Leandro Martínez, Ricardo Andrade, Ernesto G Birgin, and José Mario Martínez. Packmol: A package for building initial configurations for molecular dynamics simulations. Journal of computational chemistry, 30(13):2157–2164, 2009.

35 Olgun Guvench, Sairam S Mallajosyula, E Prabhu Raman, Elizabeth Hatcher, Kenno Vanommeslaeghe, Theresa J Foster, Francis W Jamison, and Alexander D MacKerell Jr. Charmm additive all-atom force field for carbohydrate derivatives and its utility in polysaccharide and carbohydrate–protein modeling. Journal of chemical theory and computation, 7(10):3162–3180, 2011.

36 Paul Smith, Robert M Ziolek, Elena Gazzarrini, Dylan M Owen, and Christian D Lorenz. On the interaction of hyaluronic acid with synovial fluid lipid membranes. Physical Chemistry Chemical Physics, 21(19):9845–9857, 2019.

37 David Van Der Spoel, Erik Lindahl, Berk Hess, Gerrit Groenhof, Alan E Mark, and Herman JC Berendsen. Gromacs: fast, flexible, and free. Journal of computational chemistry, 26(16):1701–1718, 2005.

38 Marco G Mazza, Nicolas Giovambattista, Francis W Starr, and H Eugene Stanley. Relation between rotational and translational dynamic heterogeneities in water. Physical review letters, 96(5):057803, 2006.

39 Leon B Lucy. An iterative technique for the rectification of observed distributions. Astronomical Journal, Vol. 79, p. 745 (1974), 79:745, 1974.

40 Lorenzo Agosta, Mikhail Dzugutov, and Kersti Hermansson. Supercooled liquid-like dynamics in water near a fully hydrated titania surface: Decoupling of rotational and translational diffusion. The Journal of Chemical Physics, 154(9), 2021.

41 Suman Dutta and J Chakrabarti. Transient dynamical responses of a charged binary colloid in an electric field. Soft Matter, 14(22):4477–4482, 2018.

42 Pinaki Chaudhuri, Ludovic Berthier, and Walter Kob. Universal nature of particle displacements close to glass and jamming transitions. Physical review letters, 99(6):060604, 2007.

