## Supplementary Information for "Heterogeneous water dynamics in the Hyaluronan-DPPC Interfaces"

### Supplementary Information:
### Heterogeneous water dynamics in the Hyaluronan-DPPC Interfaces

Anirban Paul

*Department of Physics of Complex Systems,*

*S. N. Bose National Centre for Basic Sciences,*

*Block-JD, Sector-III, Salt Lake, Kolkata 700106, India*

Jaydeb Chakrabarti

*Department of Physics of Complex Systems,*

*S. N. Bose National Centre for Basic Sciences, Block-JD,*

*Sector-III, Salt Lake, Kolkata 700106, India and*

*Department of Chemical and Biological Sciences and the Technical Research Centre,*

*S. N. Bose National Centre for Basic Sciences,*

*Block-JD, Sector-III, Salt Lake, Kolkata 700106, India*

(*)

(Dated: September 24, 2025)

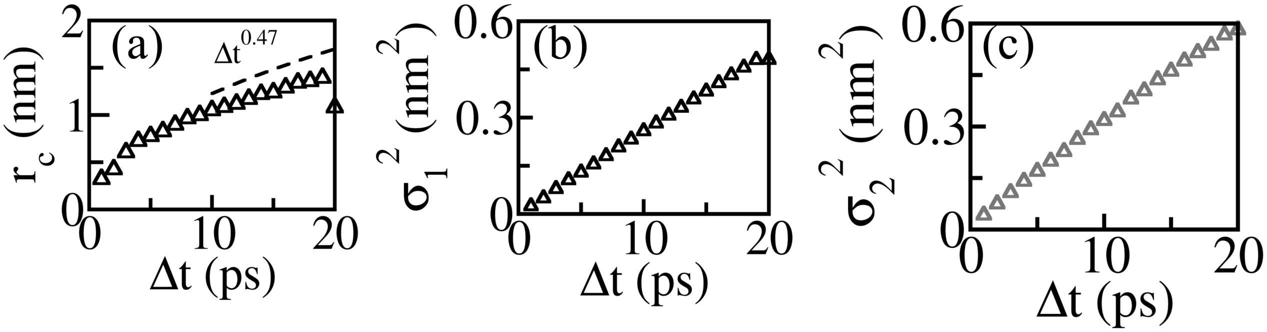


FIG. S1. (a) Variation of crossover length $r_c$ of the $G_T(r,\Delta t)$ with time $\Delta t$ for HA free case in the diffusive interface ($n_{H5}$=0). Time dependence of the widths of the two Gaussians: (b) $\sigma_1^2$ and (c) $\sigma_2^2$ of $G_T(r,\Delta t)$ for $n_{HA5}$=0.

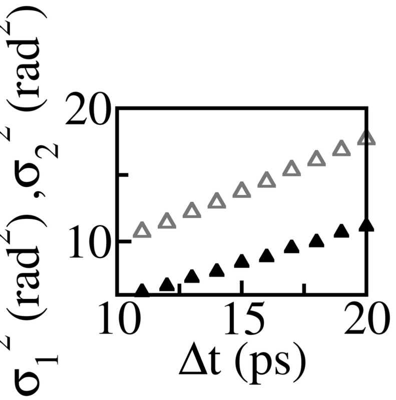


FIG. S2. Time dependence of (b) $\sigma_1^2$ (*black*) and (c) $\sigma_2^2$ (*gray*) of rotational self-vHf $G_R(\phi,\Delta t)$ in the diffusive interface for $n_{HA5}$=0.

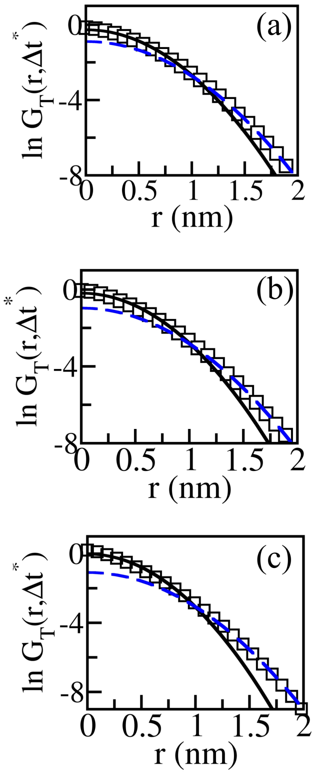


FIG. S3. $\ln G_T(r, \Delta t^*)$ vs r plot for (a) $n_{HA5}$= 10, (b) $n_{HA5}$= 30, and (c) $n_{HA5}$= 50 at $\Delta t^* = 2.0$ in the diffusive interface. Solid *black* line implies fitted central Gaussian, broken *blue* line shows fitted Gaussian tail.

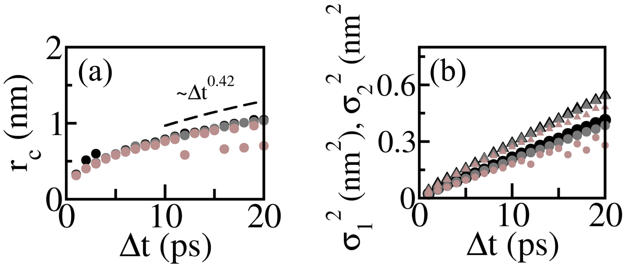


FIG. S4. (a) Variation of $r_c$ with time $\Delta t$ in the diffusive interface for different $n_{HA5}$. (b) $\sigma_1^2$ vs $\Delta t$ (*circles*) and $\sigma_2^2$ vs $\Delta t$ (*triangles*) for different $n_{HA5}$. *Black*, *gray*, and *Brown* symbols imply $n_{HA5}$=10,30 and 50 respectively.

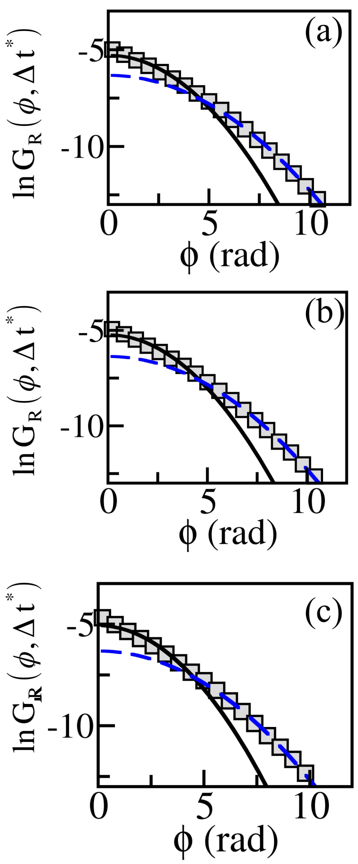


FIG. S5. $\ln G_R(\phi, \Delta t^*)$ vs $\phi$ plot for (a) $n_{HA5}$= 10, (b) $n_{HA5}$= 30, and (c) $n_{HA5}$= 50 at $\Delta t^* = 2.0$ in the diffusive interface. Solid black line implies fitted central Gaussian, broken blue line shows fitted Gaussian tail.

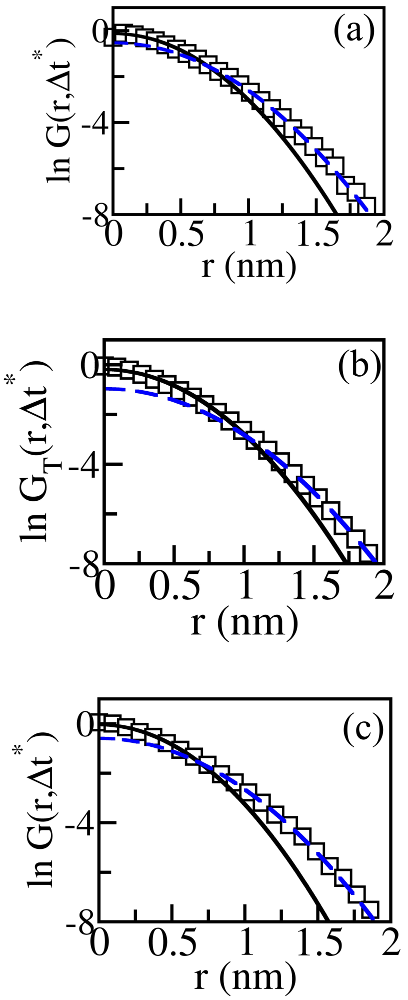


FIG. S6. $\ln G_T(r,\Delta t^*)$ vs r plot for (a) N=1, (b) N=5, and (c) N=10 at $\Delta t^* = 2.0$ in the diffusive interface. *Solid* black line implies fitted central Gaussian, *broken* blue line shows fitted Gaussian tail.

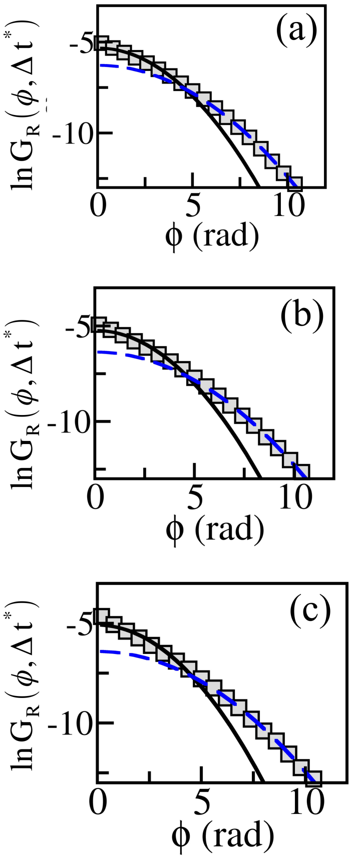


FIG. S7. $\ln G_R(\phi, \Delta t^*)$ vs $\phi$ plot for (a) N=1, (b) N=5, and (c) N=10 at $\Delta t^* = 2.0$ in the diffusive interface. *Solid* black line implies fitted central Gaussian, *broken* blue line shows fitted Gaussian tail.

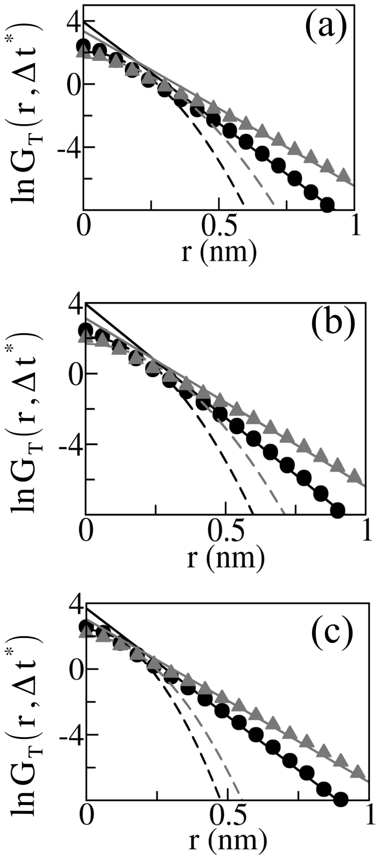


FIG. S8. $\ln G_T(r,\Delta t^*)$ vs r plot for (a) $n_{HA5}$= 10, (b) $n_{HA5}$= 30, and (c) $n_{HA5}$= 50 at $\Delta t^* = 0.5$ (*circles*) and $\Delta t^* = 1.0$ (*triangles*) in the subdiffusive hydration layer. *Broken* and *solid* lines show the fitted central Gaussian and fitted exponential tail, respectively.

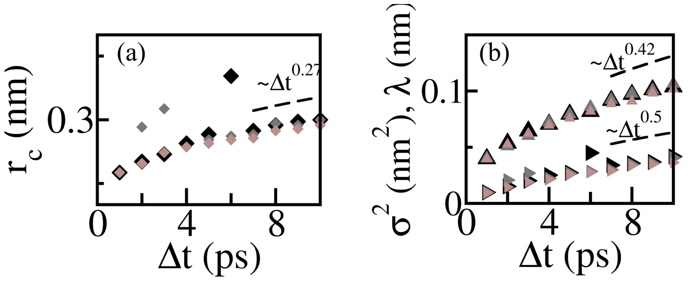


FIG. S9. (a) Variation of $r_c$ of $G_T(r,\Delta t)$ with time $\Delta t$ for different $n_{HA5}$ in the hydration layer. (b) $\sigma^2$ vs $\Delta t$ (►) and $\lambda$ vs $\Delta t$ (▲) for different $n_{HA5}$. *Black*, *gray*, and *Brown* symbols imply $n_{HA5}$=10,30 and 50 respectively.

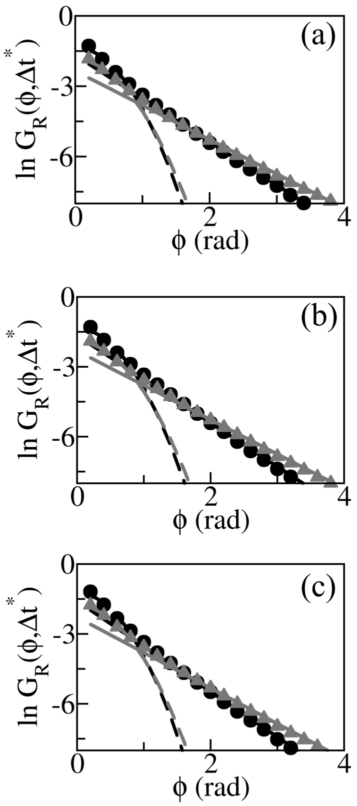


FIG. S10. $\ln G_R(\phi, \Delta t^*)$ vs $\phi$ plot for (a) $n_{HA5}$= 10, (b) $n_{HA5}$= 30, and (c) $n_{HA5}$= 50 at $\Delta t^* = 0.5$ (*circles*) and $\Delta t^* = 1.0$ (*triangles*) in the subdiffusive hydration layer. *Broken* line shows central Gaussian, *solid* line implies exponential tail.

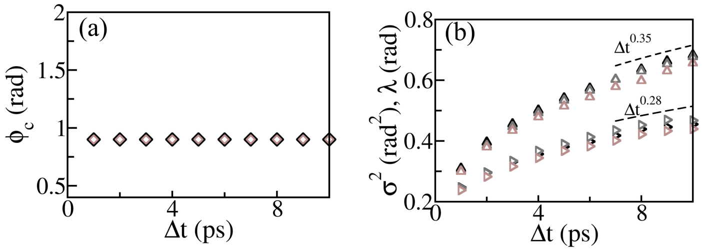


FIG. S11. (a) Variation of crossover angle $\phi_c$ of $G_R(\phi, \Delta t)$ in the hydration layer with time $\Delta t$ for different $n_{HA5}$. (b) $\sigma^2$ vs $\Delta t$ ($\triangleright$) and $\lambda$ vs $\Delta t$ ($\triangle$) of of $G_R(\phi, \Delta t)$ for different $n_{HA5}$. *Black*, *gray*, and *Brown* symbols imply $n_{HA5}$=10, 30 and 50 respectively.

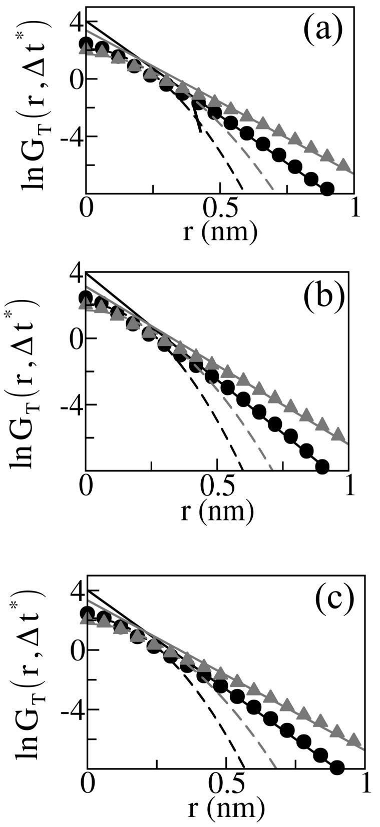


FIG. S12. $\ln G_T(r, \Delta t^*)$ vs r plot for (a) N = 1, (b) N = 5, and (c) N = 10 at $\Delta t^* = 0.5$ (*circles*) and $\Delta t^* = 1.0$ (*triangles*) in the subdiffusive hydration layer. *Broken* line shows central Gaussian, *solid* line implies exponential tail.

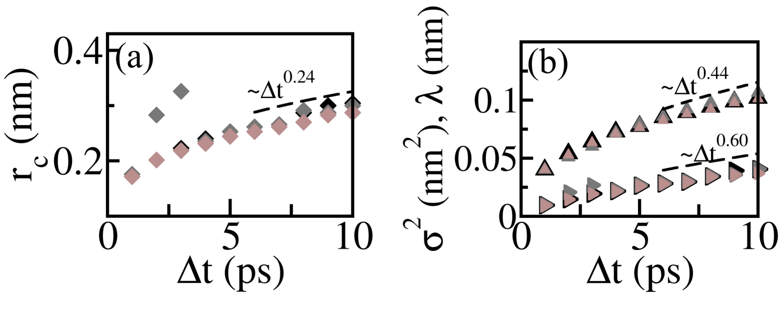


FIG. S13. The crossover length $r_c$ of $G_T(r, \Delta t)$ vs time $\Delta t$ for different N in the hydration layer. (b) $\sigma^2$ vs $\Delta t$ (►) and $\lambda$ vs $\Delta t$ (▲) for different N. *Black*, *gray*, and *Brown* symbols imply N=1,5 and 10 respectively.

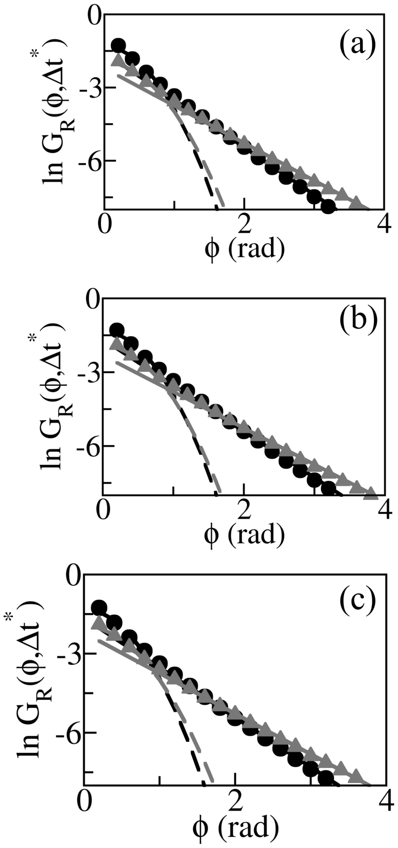


FIG. S14. $\ln G_R(\phi,\Delta t^*)$ vs r plot for (a) N = 1, (b) N = 5, and (c) N = 10 at $\Delta t^* = 0.5$ (*circles*) and $\Delta t^* = 1.0$ (*triangles*) in the subdiffusive hydration layer. *Broken* line shows Gaussian tail, *solid* line implies exponential tail.

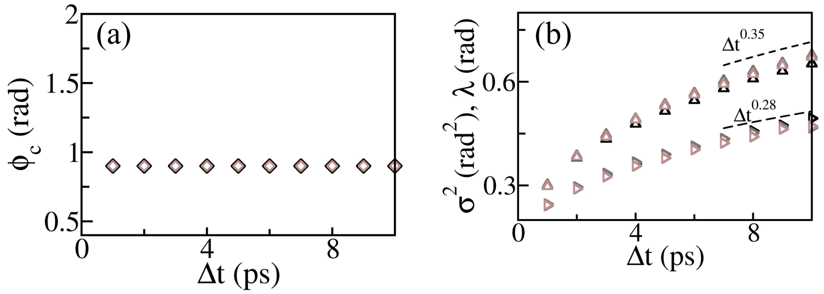


FIG. S15. Crossover angle $\phi_c$ of $G_R(\phi,\Delta t)$ vs time $\Delta t$ for different N in the hydration layer. (b) $\sigma^2$ vs $\Delta t$ ($\triangleright$) and $\lambda$ vs $\Delta t$ ($\triangle$) for different N. *Black*, *gray*, and *Brown* symbols imply N=1,5 and 10 respectively.
